# CONDITIONALLY MUTANT ANIMAL MODEL FOR INVESTIGATING THE INVASIVE TROPHOBLAST CELL LINEAGE

**DOI:** 10.1101/2023.08.02.551740

**Authors:** Khursheed Iqbal, Brandon Nixon, Benjamin Crnkovich, Esteban M. Dominguez, Ayelen Moreno-Irusta, Regan L. Scott, Ha T.H. Vu, Geetu Tuteja, Jay L. Vivian, Michael J. Soares

## Abstract

Placental development involves coordinated expansion and differentiation of trophoblast cell lineages possessing specialized functions. Among the differentiated trophoblast cell lineages are invasive trophoblast cells, which exit the placenta and invade into the uterus where they restructure the uterine parenchyma and facilitate remodeling of uterine spiral arteries. The rat exhibits deep intrauterine trophoblast cell invasion, a feature shared with human placentation, and is also amenable to gene manipulation using genome editing techniques. In this investigation, we generated a conditional rat model targeting the invasive trophoblast cell lineage. Prolactin family 7, subfamily b, member 1 (***Prl7b1***) is uniquely and abundantly expressed in the rat invasive trophoblast cell lineage. Disruption of *Prl7b1* did not adversely affect placental development. We demonstrated that the *Prl7b1* locus could be effectively used to drive the expression of Cre recombinase in invasive trophoblast cells. Our rat model represents a new tool for investigating candidate genes contributing to the regulation of invasive trophoblast cells and their contributions to trophoblast-guided uterine spiral artery remodeling.

## INTRODUCTION

The placenta creates the environment in which the fetus develops (**Maltepe and Fisher, 2015; Burton et al., 2016**). Two main functions are ascribed to the placenta and trophoblast cells, its main cellular constituents: modification of the maternal environment to support pregnancy and a role as a barrier regulating nutrient/waste flow to and from the fetus (**Soares et al., 2018; Knofler et al., 2019**). Trophoblast cells are specialized to achieve these important tasks. Hemochorial placentas, as seen in the human and some other species including the mouse and rat, possess trophoblast cell specializations that facilitate their entry into the uterine parenchyma and restructuring of uterine vasculature, which permits maternal blood to directly interface with the trophoblast cell barrier (**Pijnenborg et al., 2006; Pijnenborg and Vercruysse, 2010; Soares et al., 2018**). The intrauterine migratory abilities of these cells are best developed in the human and rat but not the mouse, which exhibits shallow invasion (**Ain et al., 2003; Pijnenborg and Vercruysse, 2010; Soares et al., 2012; Shukla and Soares, 2022**). In the human, these cells are referred to as extravillous trophoblast (**EVT**) cells, whereas the generic term, invasive trophoblast cells, is used to describe this cell population in the rat. Regulatory events controlling development of EVT and invasive trophoblast cell populations exhibit elements of conservation and are beginning to emerge from experimentation with human trophoblast stem cells and genetically modified rat models (**Chakraborty et al., 2016; Muto et al., 2021; Varberg et al., 2021; Kozai et al., 2023; Kuna et al., 2023; Vu et al., 2023**).

It is evident that some regulatory pathways controlling invasive trophoblast cells also contribute to earlier phases of placentation or more broadly to embryogenesis (**Hemberger et al., 2020; Scott et al., 2022; Vu et al., 2023**). Lentiviral strategies targeted to trophectoderm of the blastocyst have been developed to manipulate gene expression in mouse and rat trophoblast cell lineages and address some of these issues (**Georgiades et al., 2007; Okada et al., 2007; Lee et al., 2009**). Conditional mutagenesis has also become a mainstay for mouse placental research (**Woods et al., 2018**). Several trophoblast cell-specific regulatory sequences have been used to direct Cre recombinase to an assortment of different mouse trophoblast cell lineages (**Wenzel and Leone, 2007; Hu and Cross, 2011; Mould et al., 2012; Ouseph et al., 2012; Zhou et al., 2012; Crish et al., 2013; Pimeisl et al., 2013; Nadeau and Charron, 2014; Outhwaite et al., 2015; Kong et al., 2018; Wattez et al., 2019; Ozguldez et al., 2020**). Research with the rat has lagged, and model systems for the generation of conditional mutations within the rat trophoblast cell lineage are yet to be reported.

In this research project, we utilized the prolactin family 7, subfamily b, member 1 (***Prl7b1***) locus as a host for Cre recombinase. *Prl7b1* expression within the placentation site is mapped, the consequences of a *Prl7b1* null mutation described, and invasive trophoblast cell-specific actions of a *Prl7b1-Cre* recombinase rat model defined.

## RESULTS

The rat placentation site is arranged into three well-defined compartments: labyrinth zone, junctional zone, and uterine-placental interface (**Shukla and Soares, 2022; Fig. 1A**). The labyrinth zone is located at the placental-fetal interface adjacent to the junctional zone, which borders the uterine parenchyma. As gestation progresses, invasive trophoblast cells exit the junctional zone and infiltrate the uterine parenchyma, establishing a structure we define as the uterine-placental interface, which has also been called the metrial gland.

**Fig. 1.**
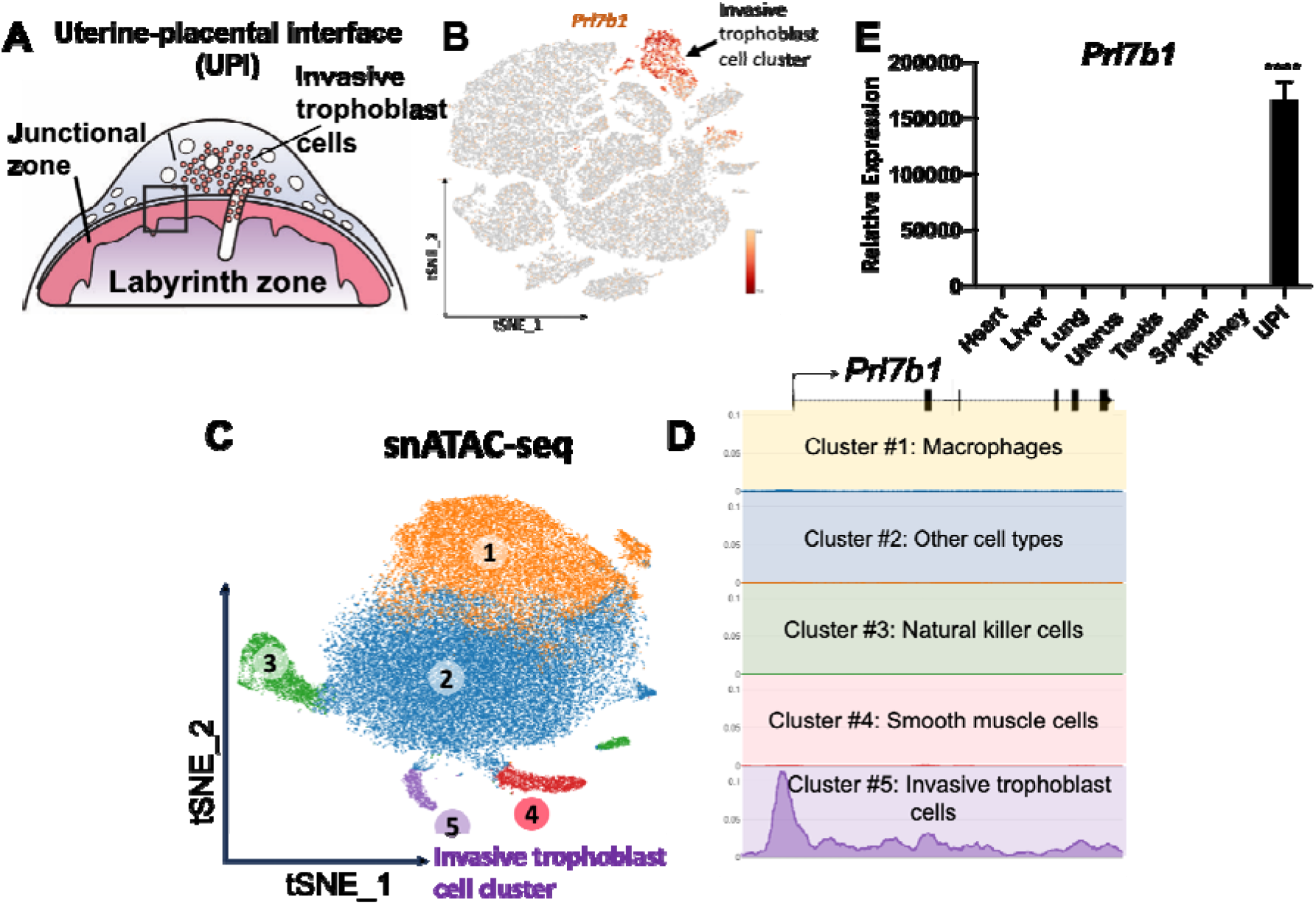
*Prl7b1* expression is specific to invasive trophoblast cells. **A**) Schematic depicting the late gestation rat placentation site. Invaded trophoblast cells are present in the uterine-placental interface (**UPI**). **B**) tSNE plot showing cell clustering on gestation day (**gd**) 19.5 UPI tissue samples, including the invasive trophoblast cell cluster (reanalyzed from **Scott et al., 2022**). **C**) t-SNE projection of single nuclei isolated from gd 19.5 UPI tissue samples and processed for single nucleus Assay for Transposase-Accessible Chromatin-sequencing (**snATAC-seq**) (reanalyzed from **Vu et al., 2023**). **D**) Chromatin accessibility profile of the rat *Prl7b1* promoter. The regulatory region upstream of the *Prl7b1* locus is highly accessible in the invasive trophoblast cell cluster in contrast to other clusters. Each colored box indicates snATAC-seq profile of each cell cluster with a corresponding color. **E**) Relative expression of *Prl7b1* transcript in various tissues determined by RT-qPCR. The histogram in panel E represent means ± SEM, n=5. Asterisks correspond to significant differences, one-way analysis of variance, * P<0.05, ** P<0.01, **** P<0.0001.

### *Prl7b1* expression in the rat placentation site

We recently performed single cell RNA-sequencing (**scRNA-seq**) and single nucleus assay for transposase-accessible chromatin using sequencing (**snATAC-seq**) for the rat uterine-placental interface from gestation day (**gd**) 15.5 and 19.5 (**Scott et al., 2022; Vu et al., 2023**). *Prl7b1* was abundantly and specifically expressed within invasive trophoblast cells (**Fig. 1B**). snATAC-seq revealed that the regulatory region of *Prl7b1* is accessible in the invasive trophoblast cell cluster (**Fig. 1C and 1D**). Among several potential host genes for Cre recombinase we selected the *Prl7b1* gene. Reverse transcription-quantitative polymerase chain reaction (**RT-qPCR**) measurements demonstrated abundant expression of *Prl7b1* transcripts in the uterine-placental interface versus any other tissue investigated (**Fig. 1E**). RT-qPCR quantification of UPI *Prl7b1* transcripts revealed progressive increases in expression as gestation proceeded (**Fig. 2A**). We confirmed the specificity of *Prl7b1* to invasive trophoblast cell lineage specific expression by in situ hybridization of the uterine-placental interface (**Fig. 2B and 2C**). *Prl7b1* transcripts were specifically localized to both endovascular and interstitial invasive trophoblast cells situated in the uterine-placental interface.

**Fig. 2.**
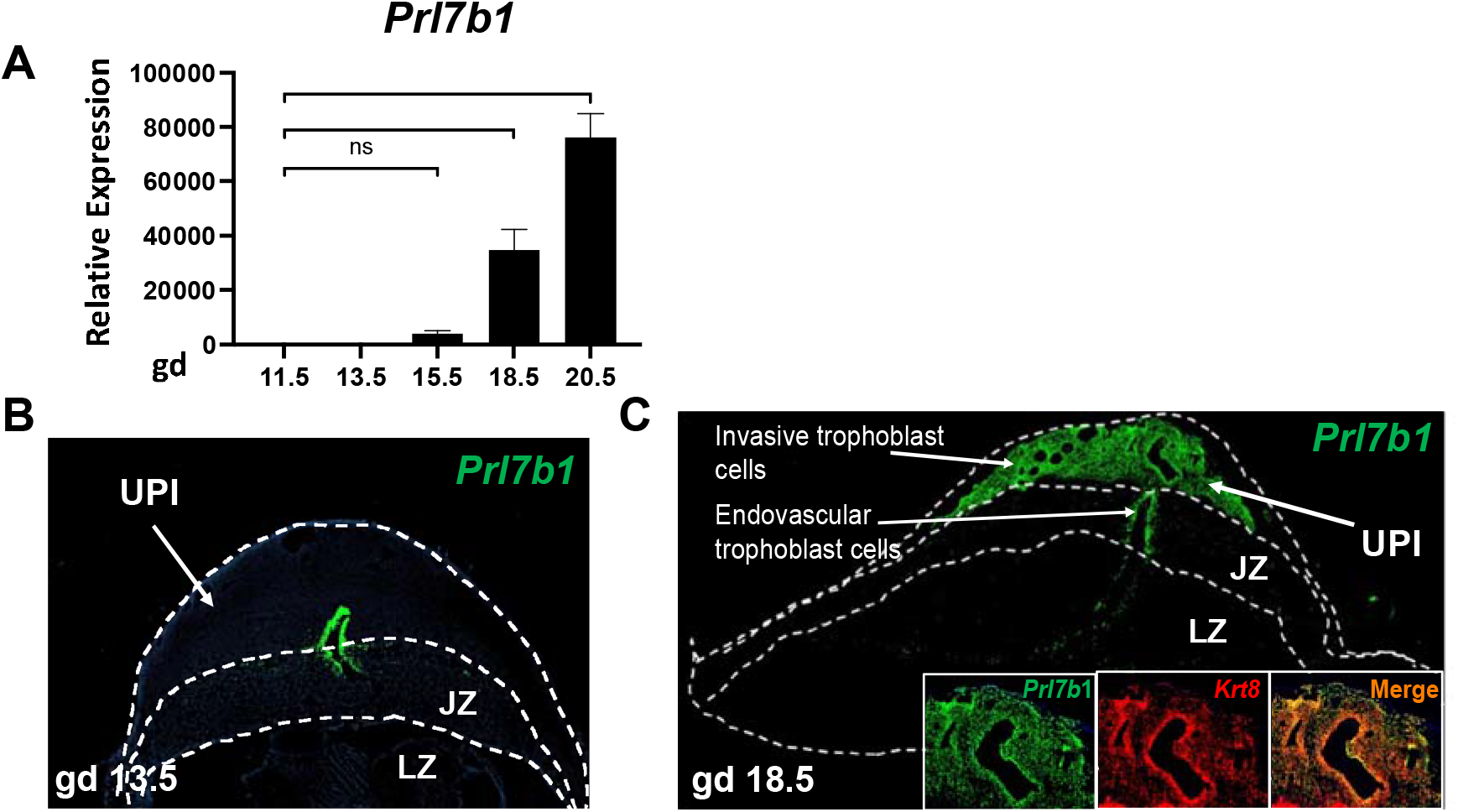
Distribution of *Prl7b1* expressing invasive trophoblast cells within the rat uterine– placental interface (UPI). **A**) Relative expression of *Prl7b1* transcripts within the UPI during gestation as measured by RT-qPCR. **B**) In situ hybridization localization of invasive trophoblast cell specific *Prl7b1* transcripts within the gd 13.5 placentation site. **C**) In situ hybridization localization of invasive trophoblast cell specific *Prl7b1* transcripts within the gd 18.5 placentation site. **D**) Co-localization of *Prl7b1* and *Krt8* transcripts within the UPI. Panel A histogram represent mean ± SEM, n = 6, 3 to 6 pregnancies. Ordinary One-way ANOVA, Dunnetts’s multiple comparisons test, ***P < 0.001, ****P < 0.0001. Labyrinth zone (**LZ**), junctional zone (**JZ**), and uterine-placental interface (**UPI**).

#### Effect of Prl7b1 disruption on fertility and pregnancy outcomes

A global *Prl7b1* deficient rat model was generated by CRISPR/Cas9 genome editing. A 272 bp deletion was generated that removed all of exon 1 and part of intron 1 (**Fig. 3A, 3B and 3C**). *Prl7b1*^Δ272^ heterozygote intercrosses generated litters of expected size and Mendelian ratio (**Table 1**). At birth, pups carrying *Prl7b1*^Δ272^ alleles (heterozygotes and homozygous nulls) were indistinguishable from wild-type littermates. Homozygous *Prl7b1*^Δ272^ rats were fertile and displayed no obvious phenotypic abnormalities. Consequently, some phenotypic comparisons proceeded on wild type intercrosses versus homozygous *Prl7b1*^Δ272^ intercrosses. Phenotypic assessments were made at gd 13.5 and 18.5 (**Fig. 3D)**. Litter size did not differ between wild type and null pregnancies (**Fig. 3D**). Additionally, the organization of gd 13.5 placentation sites was not significantly affected by PRL7B1 deficiency (**Fig. 3E**). Junctional zone and labyrinth zone compartments were well-defined in both genotypes, as was the extent of intrauterine trophoblast invasion at gd 18.5 (**Fig. 3E**). Postnatal litter size and developmental outcomes were similar among wild type intercross breeding and PRL7B1-deficient intercross breeding. Thus, PRL7B1 deficiency does not adversely affect fertility or pregnancy outcomes.

**Fig. 3.**
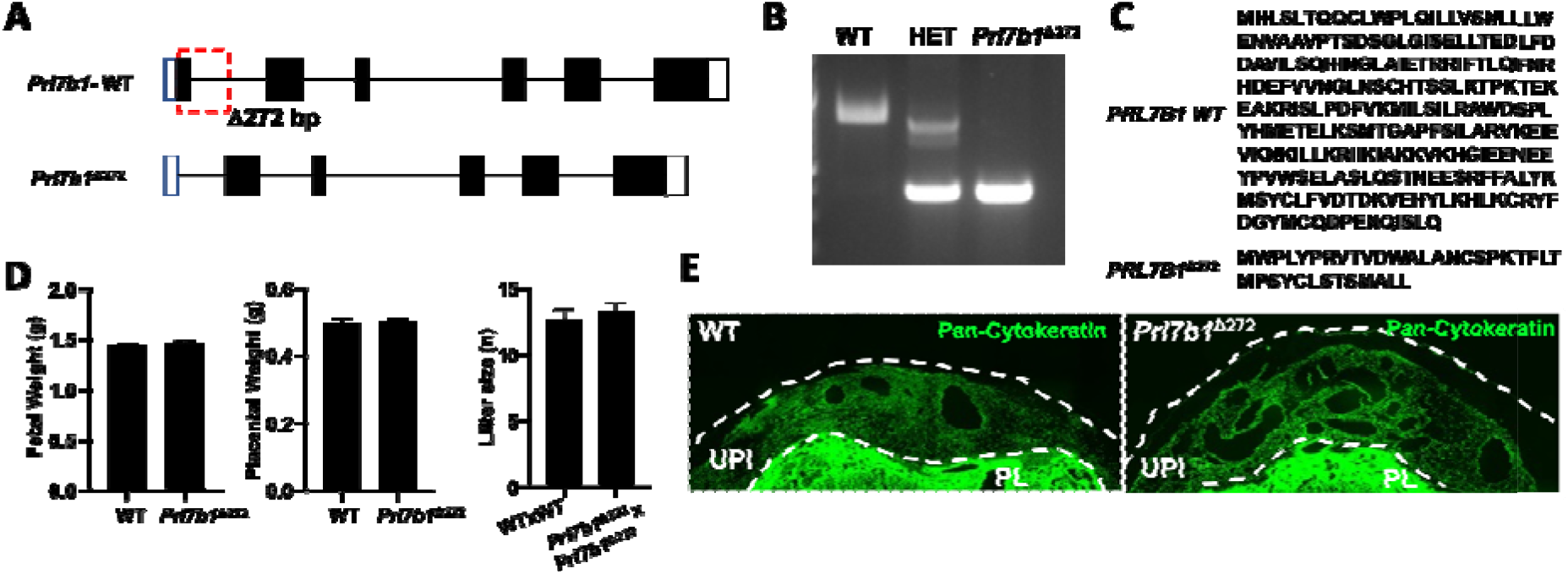
Generation of *Prl7b1* null rat and effects on pregnancy outcomes. **A**) Schematic representation of the organization of the *Prl7b1* gene, including the site for CRISPR/Cas9 mediated deletion (Δ272 bp). **B**) Genotyping of wild type and *Prl7b1* mutant alleles. Genomic DNA was isolated, PCR performed, and resolution of DNA fragments determined by agarose electrophoresis. **C**) PRL7B1 wild type and mutant amino acid sequences. **D**) Placental and fetal weights of wild type and *Prl7b1* null fetus at gd 18.5 generated from *Prl7b1*^Δ272^ heterozygote mating and **E**) litter size generated from wild type x wild type vs homozygous *Prl7b1*^Δ272^ x *Prl7b1*^Δ272^ mating. **F**) Histological structure of wild type versus *Prl7b1*^Δ272^ placentation sites. Pan-cytokeratin staining of gd 18.5 placentation sites is shown within the uterus proximal to the placenta at gd 18.5. Pan-cytokeratin positive cells within the uterus proximal to the placenta identifies invasive trophoblast cells. PL; placenta, Het; heterozygous.

#### Generation of Prl7b1-iCre knock-in

We applied CRISPR/Cas9 genome editing technique to generate *Prl7b1-iCre* knock-in rats. We inserted the codon improved Cre (**iCre**) recombinase coding sequence immediately after the start codon in Exon 1 of the *Prl7b1* locus, to generate a rat knock-in where Cre expression faithfully recapitulates the endogenous spatial and temporal *Prl7b1* expression (**Fig. 4A and 4B**). Our *Prl7b1-iCre* rat strain was designed so that the endogenous *Prl7b1* regulatory elements drive expression of Cre recombinase. Founder rats possessing the appropriate insertion of *iCre* into the *Prl7b1* gene were backcrossed to wild type rats to demonstrate successful germline transmission. Homozygous and heterozygous *Prl7b1-iCre* knock-in rats exhibited no apparent abnormalities such as embryonic development and growth or iCre toxicity. Additionally, we did not observe any pregnancy and fertility phenotypes in *Prl7b1*-Cre homozygotes (**Supplementary Fig. S1**).

**Fig. 4.**
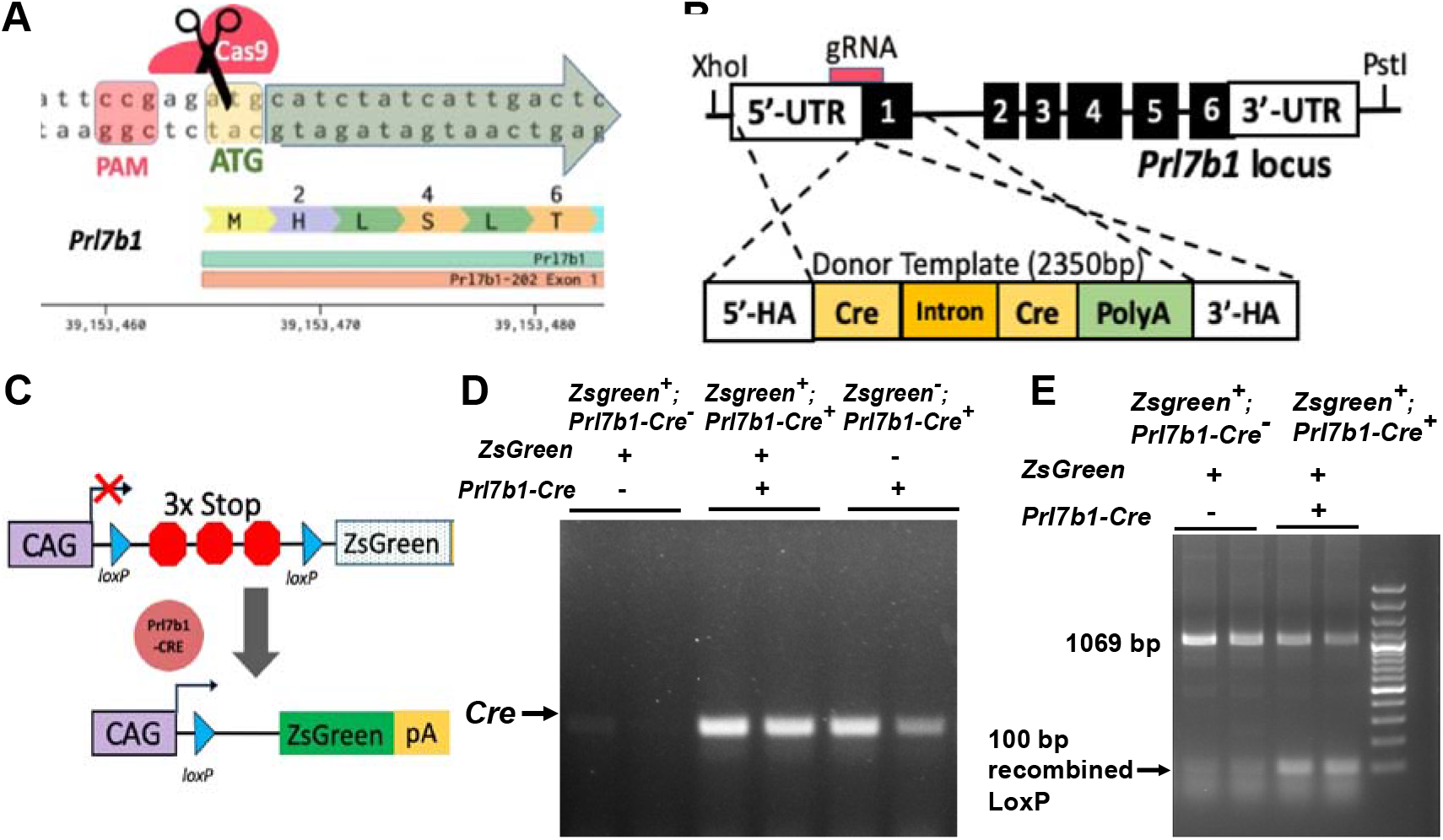
Generation of the *Prl7b1-iCre* knock-in rat driver rat strain. **A**) Schematics of CRISPR /Cas9-mediated insertion of *iCre* within the *Prl7b1* locus. **B**) Schematic illustration of Exon 1 of the wild type *Prl7b1* locus. CRISPR crRNA/tracrRNA target site is located close to *Prl7b1* ATG start codon. **C**) The reporter rat consists of ubiquitously active CAG promoter (**CAG**), LoxP-3x Stop-LoxP cassette (**LSL**) and ZsGreen (**Bryda et al., 2019**). The ZsGreen protein expresses only where *Prl7b1-iCre* is expressed. **D)** Cre transcript expression using RT-PCR in gd 18.5 UPI tissue isolated from *CAG-ZsGreen* x *Prl7b1-iCre* mating. **E)** Genotyping of gd 18.5 UPI tissue isolated from *CAG-ZsGreen* x *Prl7b1-iCre* mating showing Cre mediated loxP excision. **HA**, Homology arm; **pA**, polyA.

#### Characterization of the Prl7b1-iCre driver rat strain

To test the efficiency and specificity of Cre recombinase activity in the *Prl7b1-iCre* rat, we mated heterozygous male *Prl7b1-iCre* rats with the Cre dependent Tg(CAGloxP-STOP-loxP-ZsGreen) reporter line (**Bryda et al., 2019**). The reporter rat strain possesses a ZsGreen gene downstream of a floxed STOP cassette. When bred to a strain expressing Cre-recombinase under control of various tissue specific promoters, loxP site-specific excision of the STOP cassette occurs resulting in expression of the ZsGreen gene driven by the ubiquitously and constitutively active chicken beta-actin promoter coupled with the cytomegalovirus early enhancer (**Fig. 4C**). We initially confirmed the successful functioning of *Prl7b1-iCre* system by assessing expression of the iCre cDNA using RT-qPCR measurements of uterine-placental interface tissues (**Fig. 4D**). We confirmed the Cre recombinase-mediated recombination of loxP and site-specific excision of STOP codons in uterine-placental interface tissues by PCR (**Fig. 4E**). To examine tissue specificity of the *Prl7b1-iCre* rat line, we isolated uterine-placental interface tissues and placentas from gd 18.5 pregnant rats and imaged tissue for ZsGreen fluorescence. We observed green fluorescence of ZsGreen throughout the uterine-placental interface (**Fig. 5A and 5B**). The only *Prl7b1-iCre* positive cells capable of activating ZsGreen fluorescence within the uterine-placental interface (**UPI**) are invasive trophoblast cells. Within the placenta, ZsGreen positive cells were observed in endovascular trophoblast cells lining central placental arteries and a small subset of cells situated within the junctional zone (**Fig. 6A and 6B**). The latter may be at least part of the invasive trophoblast progenitor cell population.

**Fig. 5.**
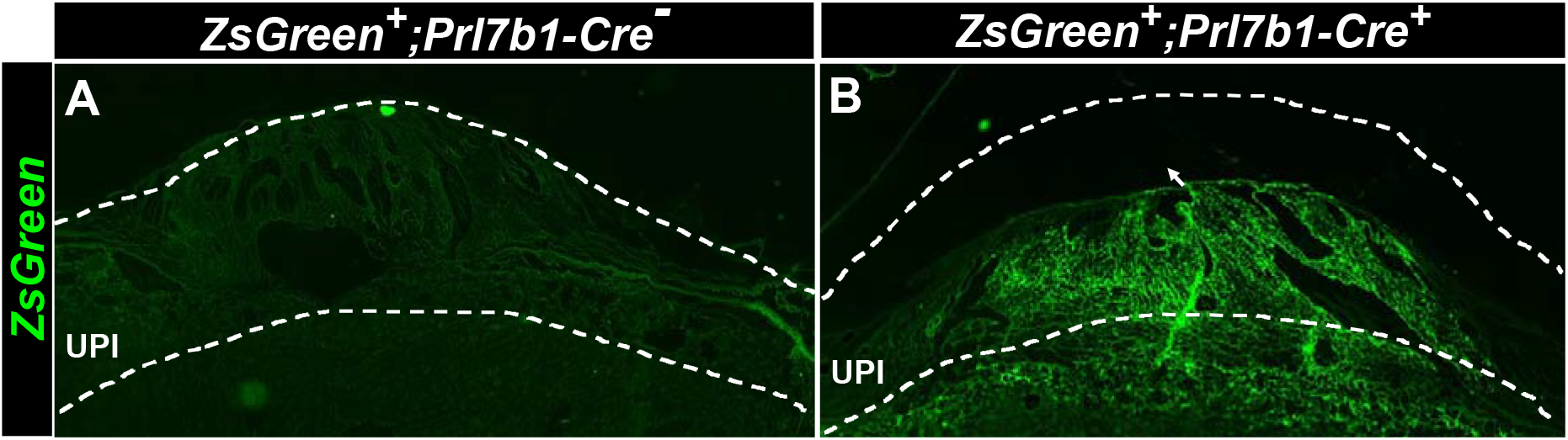
Characterization of Cre activity at the histological level in the uterine-placental interface (UPI) of the *Prl7b1-Cre* transgenic rat model. Cre recombinase mediated expression of the ZsGreen reporter gene in the *Prl7b1* expressing invasive trophoblast cells in the UPI at gd 18.5. **A)** ZsGreen fluorescence is not observed in *Zsgreen+;Prl7b1-iCre^−^* due to the absence of Cre activity in the ZsGreen UPI. **B)** ZsGreen fluorescence is specifically localized to the UPI due to *Prl7b1-Cre* recombinase mediated recombination of the ZsGreen reporter gene. <u>Arrow</u> showing the ZsGreen fluorescence in invasive endovascular trophoblast cells.

**Fig. 6.**
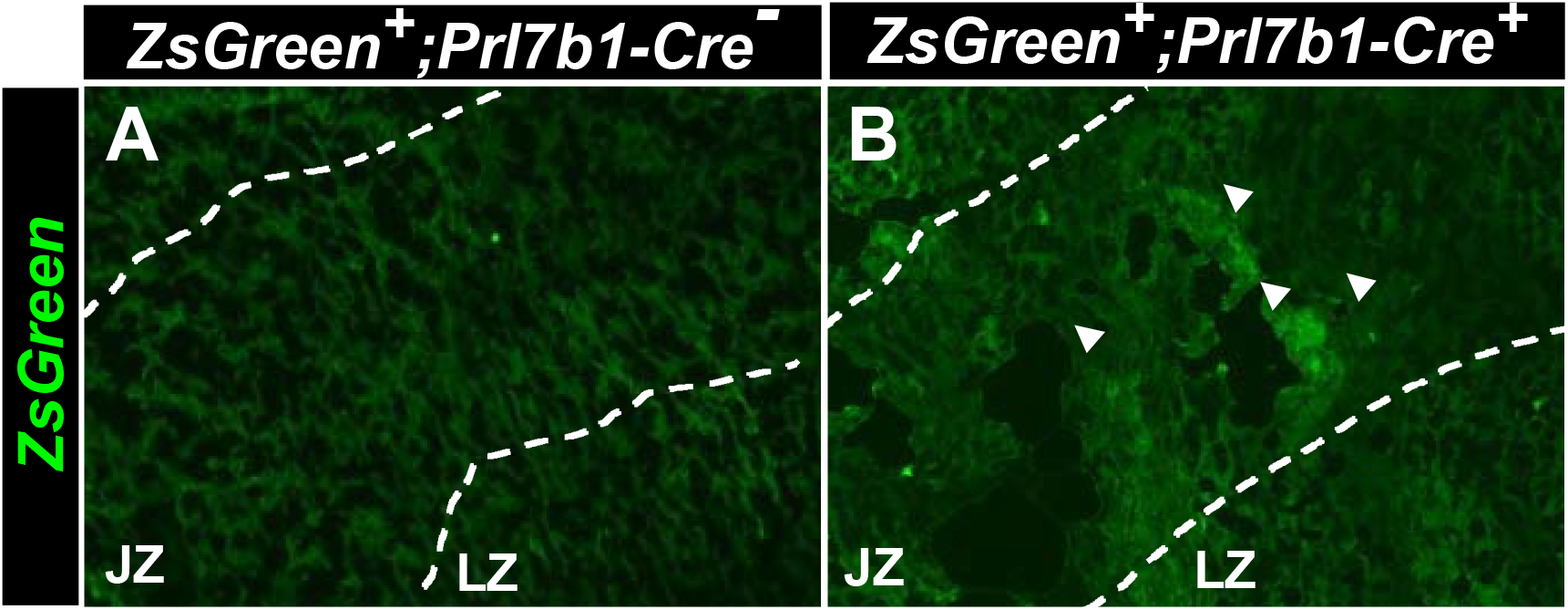
*Prl7b1* driven Cre mediated recombination during development in placental tissue. **A)** ZsGreen fluorescence is not observed due to the absence of Cre activity in ZsGreen placenta. **B)** Punctate ZsGreen fluorescence is observed in the junctional zone (<u>arrowheads</u>).

## DISCUSSION

Invasive trophoblast cells transform the uterus facilitating blood flow to the placenta and fetus (**Red-Horse et al., 2004; Velicky et al., 2016**). Disruption of trophoblast cell invasion and remodeling of uterine vasculature have been associated with obstetrical complications, including early pregnancy loss, preeclampsia, intrauterine growth restriction, and preterm birth (**Kaufmann et al., 2003; Brosens et al., 2019**). Thus, there is merit in understanding regulatory mechanisms controlling the differentiation and function of the invasive trophoblast cell lineage. In this report, we established a rat model specifically expressing Cre recombinase in invasive trophoblast cells. Cre recombinase was incorporated into the *Prl7b1* locus using CRISPR/Cas9 genome editing and specifically activated in invasive trophoblast cells situated within the uterine-placental interface.

The *Prl7b1* locus was selected as a host for *Cre* recombinase for two important reasons: i) *Prl7bl* was expressed in an invasive trophoblast-specific pattern; and ii) disruption of the *Prl7b1* does not undermine placentation or pregnancy. PRL7B1 is a member of the expanded prolactin family and includes orthologs in the rat and mouse but not human (**Soares, 2004; Soares et al., 2007**). Both endovascular and interstitial invasive trophoblast cells situated within the uterine-placental interface express *Prl7b1* (**Wiemers et al., 2003; Scott et al., 2022; Vu et al., 2023**). The absence of a detectable pregnancy-related phenotype in the *Prl7b1* mutant rat is consistent with experimentation with the *Prl7b1* mutant mouse (**Bu et al., 2016**). In the mouse, *Tpbpa* and *Prmd1* genes have been used to target Cre recombinase to trophoblast cells, including invasive trophoblast cells (**Hu and Cross, 2011; Mould et al., 2012**); however, neither locus directs expression exclusive to the invasive trophoblast cell lineage in the mouse or rat (**Ain et al., 2003; Scott et al., 2022**). The rat *Prl7b1* locus is optimal for directing mutations specifically to the invasive trophoblast cell lineage.

Effective tools for dissecting regulatory pathways controlling invasive trophoblast cells are limited. There is a plethora of experimental work performed in vitro with the goal of elucidating mechanisms underlying invasive trophoblast cell differentiation and function. Much of this work is of limited value due to inherent problems in using transformed or immortalized cell culture systems or because of the artificial nature of all in vitro analyses (**Lee et al., 2016; Soares et al., 2018**). Trophoblast stem cell models have greatly advanced the field and are best for generating hypotheses governing molecular mechanisms that can be tested in vivo (**Tanaka et al., 1998; Asanoma et al., 2011; Okae et al., 2018**). Conservation at structural and molecular levels within the human and rat uterine-placental interface are evident (**Pijnenborg and Vercruysse, 2010; Soares et al., 2012; Scott et al., 2022; Shukla and Soares, 2022; Vu et al., 2023**). Global gene disruption in the rat has proven effective in gaining insights regarding the biology of some genes involved in regulating the invasive trophoblast cell lineage (**Chakraborty et al., 2016; Muto et al., 2021; Varberg et al., 2021; Kozai et al., 2023; Kuna et al., 2023)**; however, it is apparent that genes potentially involved in controlling the biology of invasive trophoblast cells are also used in earlier phases of trophoblast cell development or in other aspects of embryonic or extraembryonic development (**Scott et al., 2022; Vu et al., 2023**). The *Prl7b1-Cre* rat model provides a valuable tool for in vivo testing of hypotheses proposed to explain mechanisms controlling the biology of the invasive trophoblast cell lineage.

## MATERIALS AND METHODS

### Animals and tissue collection

Holtzman Sprague–Dawley rats were purchased from Envigo and maintained under specific pathogen-free conditions in an Association for Assessment and Accreditation of Laboratory Animal Care–accredited animal facility at the University of Kansas Medical Center (**KUMC**). Rats were fed standard rat chow and water ad libitum and maintained in a 14-h light:10-h dark photoperiod (lights on at 0600 h). Time-mated pregnancies were established by co-housing adult female rats (8–12 weeks of age) with adult male rats (>10 weeks of age). Detection of sperm or a seminal plug in the vagina was designated gd 0.5. Pseudopregnant females were generated by co-housing adult female rats (8–12 weeks of age) with adult vasectomized male rats (>10 weeks of age). At the time of euthanasia, litter sizes and the viability of conceptuses were recorded, and tissues used for histological analysis were frozen in dry ice–cooled heptane and stored at −80 °C until processed, whereas tissues used for biochemical analyses were frozen in liquid nitrogen and stored at −80 °C until processed (**Ain et al., 2006; Chakraborty et al., 2011, 2016**). All animal procedures were approved by the KUMC Institutional Animal Care and Use Committee.

### Reanalysis of scRNA-seq and snATAC-seq

Single-cell RNA-seq data from the rat uterine-placental interface tissues (**Scott et al., 2022**) was downloaded and reanalyzed using Cellranger analysis pipelines (10x Genomics). Briefly, cellranger count was used to perform alignment, filtering, barcode counting, and UMI counting alignment. Output from multiple samples was combined using cellranger aggr (version 7.0.1). Single-nuclei ATAC-seq data from uterine-placental tissues (**Vu et al., 2023**) was downloaded and reanalyzed using cellranger-atac analysis pipelines (10x Genomics). Briefly, cellranger-atac count was used for detection of accessible chromatin peaks, count matrix generation for peaks and transcription factors. Output from multiple samples was combined using cellranger-atac aggr (version 1.1.0). scRNA-seq and snATAC-seq data was visualized using Loupe browser (10x genomics).

### Transcript analysis

Total RNA was extracted from tissues using TRIzol reagent (AM9738, Thermo Fisher). cDNAs were synthesized from total RNA (1 μg) for each sample using SuperScript 2 reverse transcriptase (18064014, Thermo Fisher), diluted ten times with water, and subjected to RT-qPCR to estimate mRNA levels. RT-qPCR primer sequences are presented in **Supplemental Table 1**. Real-time PCR amplification of cDNAs was carried out in a reaction mixture (20 μL) containing SYBR GREEN PCR Master Mix (4309155, Applied Biosystems) and primers (250 nM each). Amplification and fluorescence detection were carried out using the ABI QuantStudio PCR system (Applied Biosystems). The delta–delta Ct method was used for relative quantification of the amount of mRNA for each sample normalized to 18S RNA.

### In situ hybridization

Distributions of transcripts for *Prl7b1* and *Krt8* were determined on cryosections of rat placentation sites at various gestation days. RNAScope Multiplex Fluorescent Reagent Kit version 2 (Advanced Cell Diagnostics) was used for in situ hybridization analysis. Probes were prepared to detect *Prl7b1* (NM_153738.1, 860181, target region: 28-900) and *Krt8* (NM_199370.1, 873041-C2, target region: 134-1,472). Fluorescence images were captured on a Nikon 80i upright microscope with a Photometrics CoolSNAP-ES monochrome camera (Roper Scientific Inc). The images were processed using FIJI software.

### Generation of global Prl7b1 knock-out and Prl7b1-iCre knock-in rat models

The rat *Prl7b1* (ENSRNOG00000016742, NM_153738) gene is situated on Chromosome 17 (Chr17: 39,153,434-39,161,643) among members of the prolactin (**PRL**) family. Mutations at the *Prl7b1* locus were generated using genome editing as previously described by our laboratory (**Iqbal et al., 2021a,b; Kozai et al., 2021, 2023; Muto et al., 2021; Varberg et al., 2021; Kuna et al., 2023**) with some modifications. In brief, 4- to 5-week-old donor rats were intraperitoneally injected with 30 units of equine chorionic gonadotropin (G4877, Sigma-Aldrich), followed by an intraperitoneal injection of 30 units of human chorionic gonadotropin (C1063, Sigma-Aldrich) ∼ 46 h later, and immediately mated with adult male rats. Zygotes were flushed from oviducts the next morning (gd 0.5) and maintained in M2 medium (MR-015-D, EMD Millipore) supplemented with bovine serum albumin (A9647, Sigma-Aldrich) at 37 °C in 5% CO_2_ for 2 h. CRISPR RNA (**crRNA**) sequence was designed to target Exon 1 (AGTCAATGATAGATGCATCTCGG) and near the translation start codon (**ATG**) for the rat *Prl7b1* gene (NM_153738). crRNAs were annealed with tracrRNA in equimolar concentrations to generate crRNA:tracrRNA duplexes (guide RNA). The ribonucleoprotein (**RNP**) complex consisting of the Cas9 protein and a synthesized crRNA/tracrRNA along with a double-stranded deoxyribonucleic acid (**dsDNA**) donor template were microinjected into embryonic day (E) 0.5 rat zygotes at a concentration of 25 ng/μL in Tris-EDTA buffer (pH 7.4). The 2,350 bp donor template consisted of an iCre cassette with a Kozak sequence and an intron (1100 bp) followed by a heterologous poly A signal sequence (200 bp) (**Wu et al., 2008**) and two 500 bp (1000 bp) homology arms (**Fig. 4A and 4B**). We incorporated Kozak and synthetic intron elements into the iCre transgene to increase transgene expression. The donor template DNA was excised from the plasmid backbone prior to co-microinjection with the RNP. Genome editing reagents were obtained from Integrated DNA Technologies. Microinjections were performed using a Leica inverted microscope and an Eppendorf Microinjection system. Manipulated zygotes were transferred to oviducts of pseudopregnant rats (20–30 zygotes per rat). Offspring were screened for deletions and insertions of *iCre* sequence within the *Prl7b1* gene. Insertion boundaries were verified by Sanger DNA sequencing. PCR primers used for genotyping of the genetically altered rats are listed in **Supplemental Table 1**. Germline transmission of the mutated gene was determined in F1 rats by backcrossing F0 founder rats with wild type rats. Detection of a mutation in F1 rats identical to the mutation present in the parent F0 rat was considered successful germline transmission. *Prl7b1* global knock-out and *Prl7b1-Cre* knock-in models will be available through the Rat Resource & Research Center (University of Missouri, Columbia, MO; www.rrrc.us).

### Histological and immunohistochemical analyses

Immunohistochemical analyses were performed on 10-μm frozen tissue sections using indirect immunofluorescence. Primary antibodies to vimentin (1:1000, V6630, Sigma-Aldrich) and pan-cytokeratin (1:1000, F3418, Sigma-Aldrich) were used in the analyses. Goat anti-mouse IgG conjugated with Alexa 488 (1:1000 dilution, A11029, Thermo Fisher) and goat anti-mouse IgG conjugated with Alexa 568 (1:400 dilution, A11031, Thermo Fisher) were used to detect primary antibodies. Fluoromount-G™, with 4′6-diamidino-2-phenylindole (DAPI, 00-4959-52, Thermo Fisher), was used to visualize nuclei and as mounting medium. Processed tissue sections were examined, and images were captured with a Nikon Eclipse 80i upright microscope equipped with a charge-coupled device camera (Nikon).

#### Statistical analysis

Student’s *t*-test and Ordinary One-way ANOVA followed by Dunnetts’s multiple comparisons test were performed, where appropriate, to evaluate the significance of the experimental manipulations. Results were determined statistically significant when P<0.05.

## ACKNOWLEDGEMENTS

We thank Stacy Oxley and Brandi Miller for administrative assistance.

## COMPETING INTERESTS

The authors declare no competing or financial interests.

## AUTHOR CONTRIBUTIONS

K.I., J.L.V., and M.J.S. designed research; K.I., B.N., A.M., B.C., E.M.D., and R.L.S. performed research; K.I., H.T.H.V., G.T., and M.J.S. analyzed data; K.I. and M.J.S. prepared the manuscript; All authors contributed to editing the manuscript. Funding acquisition: K.I., M.J.S.

## FUNDING

This research was supported by postdoctoral fellowships from the Kansas IdeA Network of Biomedical Research Excellence (P20 GM103418 to A.M-I., E.M.D.) and the Lalor Foundation (A.M-I., E.M.D.), a National Institutes of Health (NIH) National Research Service Award (HD104495, R.L.S.) and NIH grants [HD104071 (K.I.), HD020676, HD099638, HD104033, HD105734], and the Sosland Foundation.

---

**Supplementary Fig 1.**
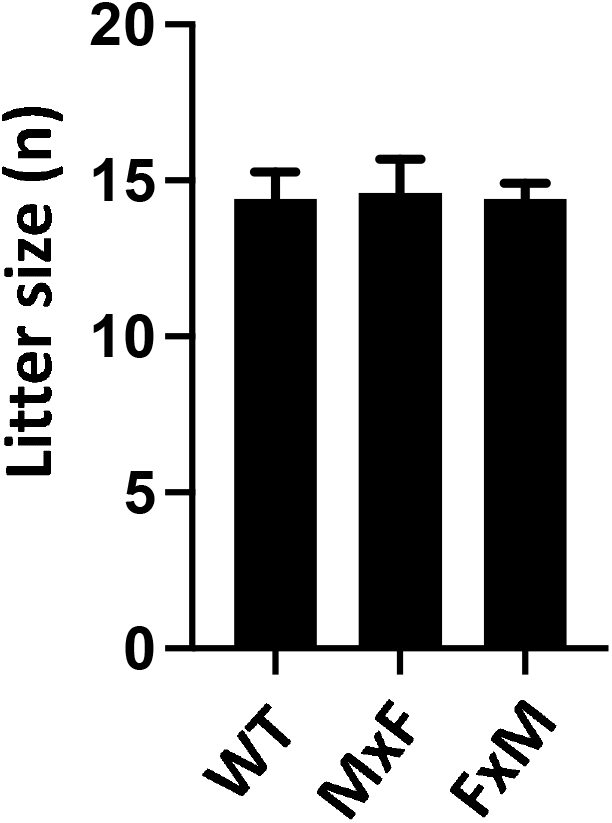
*Prl7b1-iCre* insertion does not affect litter size. Breeding homozygous *Prl7b1-iCre* males and females in different combinations (Male x Female; **M x F**, Female x Male, **F x M**) resulted in a comparable number of offspring as observed to wild-type rats. (n=5 in each group).

## Notes

### Competing Interest Statement

The authors have declared no competing interest.

